# Respiration aligns perception with neural excitability

**DOI:** 10.1101/2021.03.25.436938

**Authors:** Daniel S. Kluger, Elio Balestrieri, Niko A. Busch, Joachim Gross

**Affiliations:** Institute for Biomagnetism and Biosignal Analysis, University of Muenster, Muenster, Germany; Otto Creutzfeldt Center for Cognitive and Behavioral Neuroscience, University of Muenster, Muenster, Germany; Institute of Psychology, University of Muenster, Muenster, Germany; Centre for Cognitive Neuroimaging, Institute of Neuroscience and Psychology, University of Glasgow, Glasgow, UK

## Abstract

Recent studies from the field of interoception have highlighted the link between bodily and neural rhythms during action, perception, and cognition. The mechanisms underlying functional body-brain coupling, however, are poorly understood, as are the ways in which they modulate behaviour. We acquired respiration and human magnetoencephalography (MEG) data from a near-threshold spatial detection task to investigate the trivariate relationship between respiration, neural excitability, and performance. Respiration was found to significantly modulate perceptual sensitivity as well as posterior alpha power (8 – 13 Hz), a well-established proxy of cortical excitability. In turn, alpha suppression prior to detected vs undetected targets underscored the behavioural benefits of heightened excitability. Notably, respiration-locked excitability changes were maximised at a respiration phase lag of around - 30° and thus temporally preceded performance changes. In line with interoceptive inference accounts, these results suggest that respiration actively aligns sampling of sensory information with transient cycles of heightened excitability to facilitate performance.

## Introduction

Human respiration at rest is a continuous, rhythmic sequence of active inspiration and passive expiration ^1^. Even the active phase seems effortless, however, thanks to medullar microcircuits like the preBötzinger complex who autonomously and periodically drive respiration ^2^. Despite being largely automatic, breathing can also be adapted top-down when required, for example during speech or laughter ^3^. The bidirectionality between respiratory control and higher cognitive functions comes to no surprise, given that key structures like the preBötzinger complex (through the central medial thalamus) and the olfactory bulb are intricately connected to both the limbic system ^4^ and the neocortex ^5^.

In turn, neural oscillations in the cortex have long been established as sensitive markers of brain states ^6^, which has brought increasing attention to questions of respiration-brain coupling. While there is a rather extensive body of literature on the interaction of respiratory and neural rhythms in animals, these links are only beginning to be addressed in the human brain. Recently, both invasive ^7^ and non-invasive work ^8^ has shown respiration to modulate neural oscillations across a wide network of cortical and subcortical areas, including those not typically associated with olfaction. Moreover, modulatory effects of respiration have been demonstrated in motor ^9,10^ as well as in cognitive tasks ^7,11^. Taken together, these studies provide strong evidence for breathing-related changes in neural signalling that, particularly in task contexts, translate into changes in behaviour. Yet, fundamental questions in the trivariate relationship between respiration, neural oscillations, and behaviour remain unanswered. Respiration-locked behavioural changes have so far been demonstrated in ‘higher-level’ cognitive paradigms (e.g. visuospatial rotation or emotional judgement tasks), which complicates the identification of a clear mechanism by which respiration shapes perception and behaviour. Here, we used a simple perception task to propose neural excitability as a key moderator underlying behavioural changes coupled to respiration phase.

Cross-frequency phase-amplitude coupling is a well-established mechanism of neural information transfer across spatiotemporal scales ^12^ and provides an intuitive explanation how the link between respiration phase and oscillatory amplitudes is potentially implemented: In mice, slow respiration-induced rhythms within the olfactory bulb were shown to be transmitted through piriform cortex and subsequent cortico-limbic circuits to modulate the amplitude of faster oscillations in upstream cortical areas ^13,14^. In order to plausibly explain the perceptual and behavioural modulation effects in humans, the respiratory rhythm has to be coupled to specific neural rhythms that code transient brains states of heightened susceptibility for sensory stimulation. These phasic cycles of neural excitability determine the intensity (or *gain*) of early sensory responses in order to amplify relevant or salient stimuli at the expense of irrelevant ones ^15,16^. Excitability has been shown to vary over the respiration cycle in rats ^17^ and is tightly coupled to the amplitude of human cortical alpha oscillations (8 - 13 Hz): Particularly in the visual system, prestimulus alpha power over parieto-occipital sensors is inversely related to early visual responses ^18–20^. As a consequence, one highly replicated finding is that a prestimulus decrease in alpha power leads to increased detection rates for near-threshold stimuli ^21–23^.

Coming back to respiration-brain coupling and excitability, not only do performance levels in cognitive tasks fluctuate over the respiration cycle ^11^, but so does spontaneous alpha activity ^8^, most likely due to its dependence on arterial CO_2_ levels ^24^. Therefore, respiration-entrained fluctuations in neuronal excitability represent a promising mechanism for unifying neural and behavioural findings. Both theoretical accounts and evidence from animal and human studies strongly suggest that this link is by no means accidental, but an example of *active sensing* ^15^. While it is admittedly easier to imagine the high-frequency sniffing and whisking of mice as an active sampling of sensory stimuli, a similar case can be made for human respiration from the perspective of predictive processing ^25^: In temporally coordinating the breathing act and internal brain dynamics (i.e., heightened excitability), the sampling of bottom-up sensory information can be aligned with top-down predictive streams ^26^. This active view of respiration is corroborated by reports that human participants spontaneously inhale at trial onset in a self-paced cognitive task ^11^, effectively aligning stimulus processing with the inspiration phase. Overall, evidence from the lines of research examined so far point to respiration as a mechanism to synchronise perception and neural excitability, which in turn might influence human performance in a variety of tasks. However, this mechanism currently lacks a clear demonstration, which we aim to provide with the present experiment. We simultaneously recorded respiration, whole-head MEG, and performance measures from thirty human participants in a spatial detection task to address the questions introduced earlier: First, we investigated whether respiration cyclically affects sensitivity towards near-threshold stimuli and how this link develops over the respiration cycle. Second, using parieto-occipital alpha power as a well-established proxy of excitability, we assessed respiration-locked excitability changes and how they are related to performance. Finally, we aimed to illuminate the overarching link between respiration-excitability and respiration-performance modulations.

## Results

### Respiration modulates perception

Participants performed a simple detection task in which Gabor patches were briefly presented at near-threshold contrast either to the left or to the right of a central fixation cross. After a short delay, participants were to report via button press whether they had seen the target on the left, the right, or no target at all (see Fig. 1a and Methods for details). An adaptive staircase (QUEST) was used to adjust the contrast of target trials in a way that performance would settle at a hit rate of around μ_HR_ = .60 (Fig. 2a). Across participants, we observed an average hit rate (i.e. the proportion of detected targets) of M_HR_ = .54 ± .05 (M ± SD), which was reasonably close to the staircase’s desired hit rate (see Supplementary Fig. 1 for the distribution of individual hit rates).

**Fig. 1.**
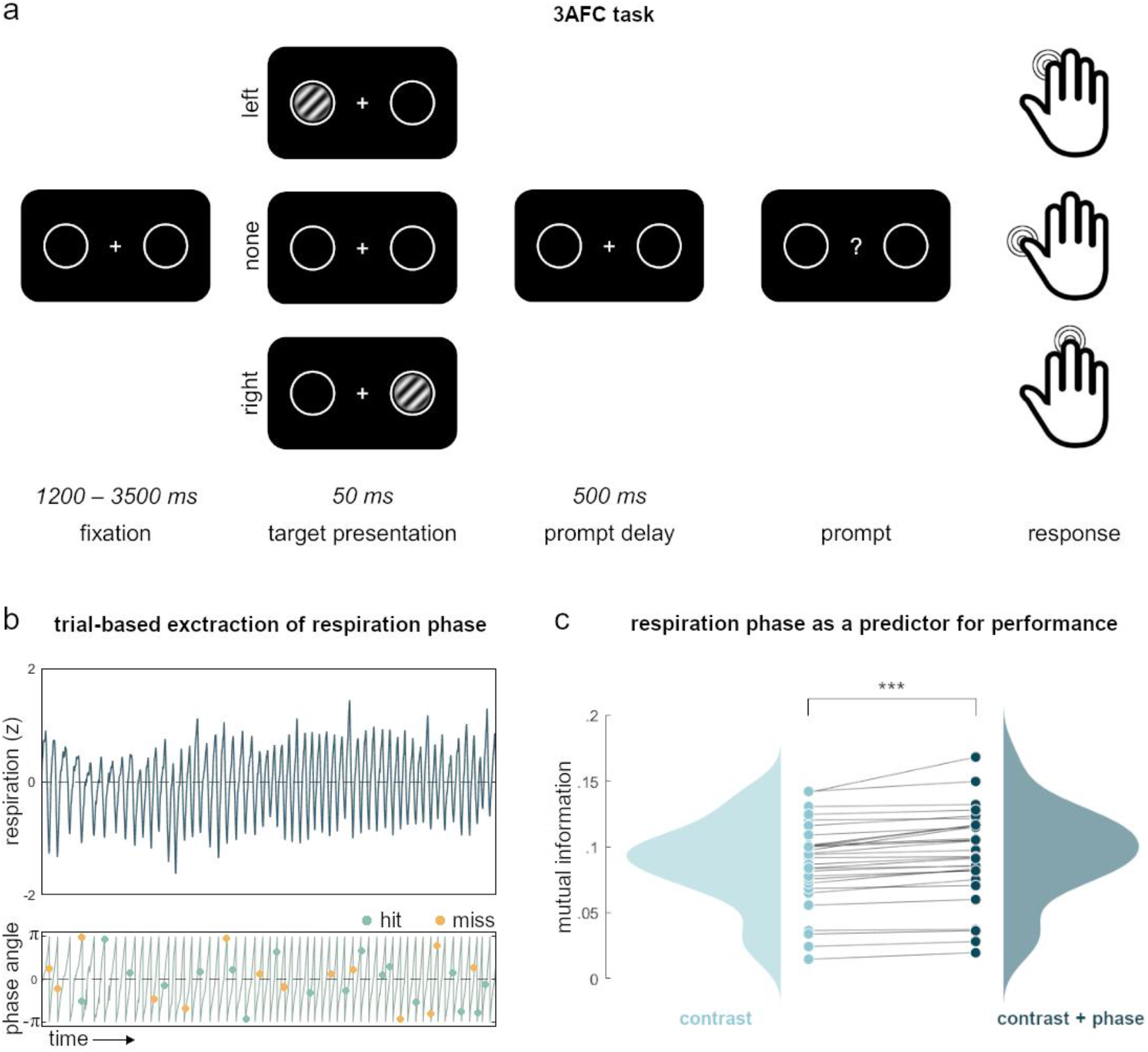
Task and behavioural results. **a**, In the experimental task, participants kept their gaze on a central fixation cross while a brief, near-threshold Gabor patch (magnified for illustrative purposes) was randomly presented either on the left or the right side of the screen. During catch trials, no stimulus was presented. After a brief delay, participants were prompted to indicate whether they saw a stimulus on the left (index finger) or on the right side (middle finger), or no stimulus at all (thumb). **b**, Exemplary segment of respiration recordings (top) plotted against its phase angle (bottom). Over time, targets were randomly presented over the respiration cycle and could be either detected (*hits*, green) or undetected (*miss*, yellow). **c**, Raincloud plot shows individual mutual information computed on the trial-level between detected/undetected and stimulus contrast (left) and detected/undetected and stimulus contrast plus respiration phase (right). Mutual information was significantly enhanced by adding respiration phase (Wilcoxon signed rank test: *z* = 4.78, *p* < .001). Raincloud plot generated with the *RainCloudPlots* toolbox for Matlab^27.^

**Fig. 2.**
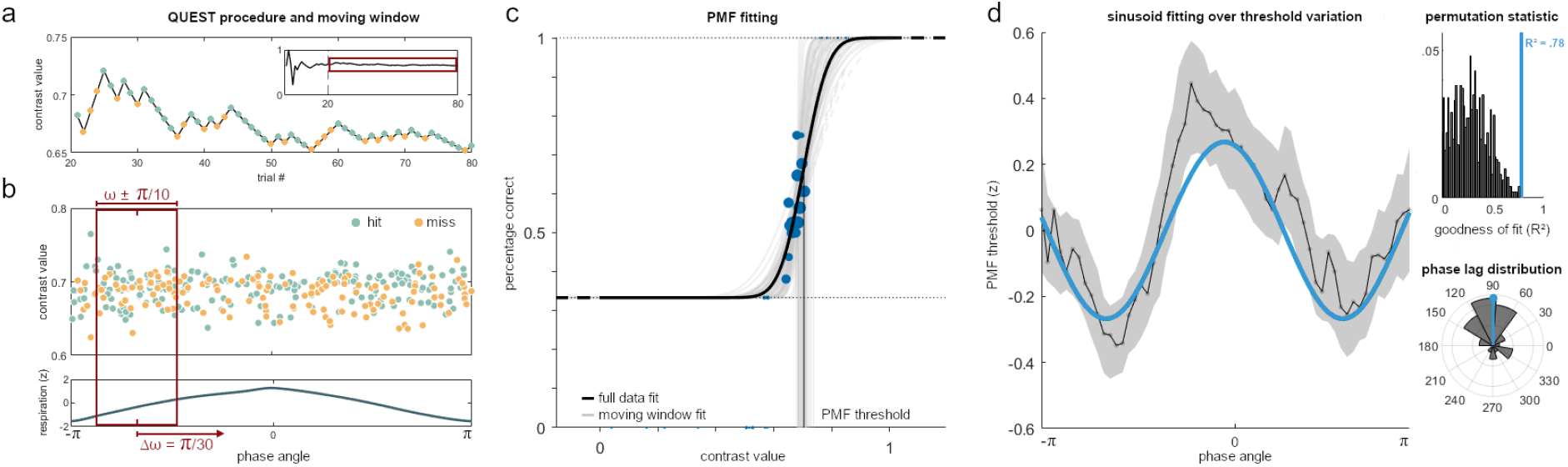
Respiration phase modulates performance. **a**, The first twenty trials of each run were discarded to allow initiation of the QUEST staircase (inset shows contrast variation of the full run). Extending the behavioural analysis shown in Fig. 1, we obtained contrast values and respiration phase angle of hits (green) and misses (orange). Catch trials did not inform the QUEST staircase and are not shown. **b**, Top panel shows the contrast values of all hit and miss trials of one exemplary participant sorted by the respiration phase angle at which the targets were presented. Moving along the respiration cycle (bottom) in increments of π/30, we applied a moving window with a width of π/5 to select a subset of trials for a phase-locked refitting of the psychometric function. **c**, The psychometric function was first fitted to the full data set (black line, shown here for one typical participant). All parameters except the threshold were then fixed and used as priors for fitting the PMF iteratively to an angle-specific subset of trials (grey lines). Blue dots mark clusters of single trials scaled by frequency of occurrence. For each participant, the procedure of trial selection and re-fitting was repeated 60 times until all trials from all respiration phase angles had been sampled. **d**, Left: The moving window approach resulted in individual courses of PMF threshold over respiration angles whose normalised mean is shown here ± SEM (black line and shading). These modulations corresponded to contrast value changes within the range of [.69 .72]. Blue line shows the best group-level sinusoid fit at 1.55x respiration frequency. Top right: The empirical goodness of fit (adjusted R^2^ = .78) was highly significant (*p* < .001), as evident from the comparison with its null distribution constructed from 1000 permutations using random shifts of the respiration signal. Bottom right: Group-level phase lag distribution of sinusoid fits.

While individual hit rates were obviously dependent on the contrast of each target, one of our main aims was to assess the independent influence of the interoceptive breathing signal. As a first analysis of potential respiration-related performance changes, we employed a model comparison of single-trial mutual information. For each participant, we first computed mutual information between the discrete factor detected/undetected and the continuous factor of Copula-normalised stimulus contrast. In a second computation, we included another continuous factor with sine and cosine of the respiration phase angle at which each trial was presented (see Fig. 1b for an illustration). On the group level, we then compared the mutual information with and without respiration phase by testing participant-level differences against zero. A Wilcoxon signed rank test confirmed that mutual information was significantly enhanced by adding respiration phase information (*z* = 4.78, *p* < .001; see Fig. 1c). Having provided first evidence for a functional link between respiration and perceptual performance, we next conducted an in-depth analysis to illustrate how such behavioural modulation occurs independently of target contrast.

### Respiration changes the psychometric function

Since the previous analysis had established a significant relationship between respiration and perceptual performance, we next addressed the question if this relationship is mediated by a respiration-induced change of the individual psychometric function that relates stimulus contrast to perceptual performance.

In order to assess respiration-related changes in perceptual accuracy irrespective of QUEST-induced changes in stimulus intensity, we exploited single-trial information about both stimulus (i.e., its contrast) and performance (i.e., whether it was detected or not) in an iterative re-fitting of the PMF: For each participant, we first fitted an overall PMF to all target trials (see Supplementary Fig. 2 for individual PMF fits). The resulting parameters (threshold, width) were used as priors for a moving window approach in which we iteratively re-fitted the psychometric function to a subset of trials presented at a certain range of respiration angles (see description above and Fig. 2b-c). For each of the 60 overlapping phase angle bins ranging from -π to π, we thus obtained a threshold estimate characterising perceptual performance at that respiration phase: With a fixed response criterion of μ_HR_ = .60, a lower threshold indicates higher accuracy, as a lower stimulus intensity was sufficient to yield the same level of performance. Indeed, individual thresholds showed systematic variations across the respiratory cycle (Fig. 2d), indicating that respiration rhythmically changes perceptual threshold. To statistically validate this finding, we fitted a sinusoid function to the normalised individual courses of threshold variation, yielding a high model fit of adjusted R^²^ = .78 (Fig. 2d). Significance on the group level was assessed by means of nonparametric permutation testing, confirming that our empirical model fit was significantly higher than the 95^th^ percentile of a null distribution constructed from 1000 random shifts of respiration phase (*p* < .001, Fig. 2d).

In order to characterise phase consistency between subjects at the best fitted frequency, we finally computed a fast Fourier transform (FFT) of individual threshold variation courses. This approach complemented the previous analysis by computing a phase metric at the single-subject level using an analytic computation that does not require numerical optimisation. For each participant, we selected the phase angle of the vector in the complex plane closest to the fitted frequency and tested for angle uniformity across subjects using a Rayleigh test. We found a significant phase alignment at a frequency of F_M_ = 1.52 cycles/T (*z* = 5.68 *p* = .003), congruent with the best fitted frequency at the population level (F_P_ = 1.55 cycles/T).

Taken together, our analysis demonstrates that the individual psychometric function changes significantly across the respiratory cycle. The next step was to assess the role of alpha oscillations and how they are linked to performance changes. If respiration, as we propose, does shape perception through changes in cortical excitability, we can form precise assumptions regarding parieto-occipital alpha power as a proxy of excitability.

### Alpha power is related to behavioural performance

For the excitability hypothesis to hold true, changes in pre- and peristimulus alpha power over parieto-occipital sensors would first have to be linked to changes in behaviour. Replicating a rich body of previous work (reviewed in ^28^), we compared parieto-occipital MEG power spectra for detected and undetected targets (see Fig. 3a). Time-frequency analyses confirmed significant suppression of alpha power prior to presentation of targets which were later detected (vs undetected, see Fig. 3a for cluster-corrected results across ROI sensors). Alpha suppression persisted as long as one second prior to target onset (Fig. 3b) and was mainly localised over occipital and posterior parietal sensors (Fig. 3c).

**Fig. 3.**
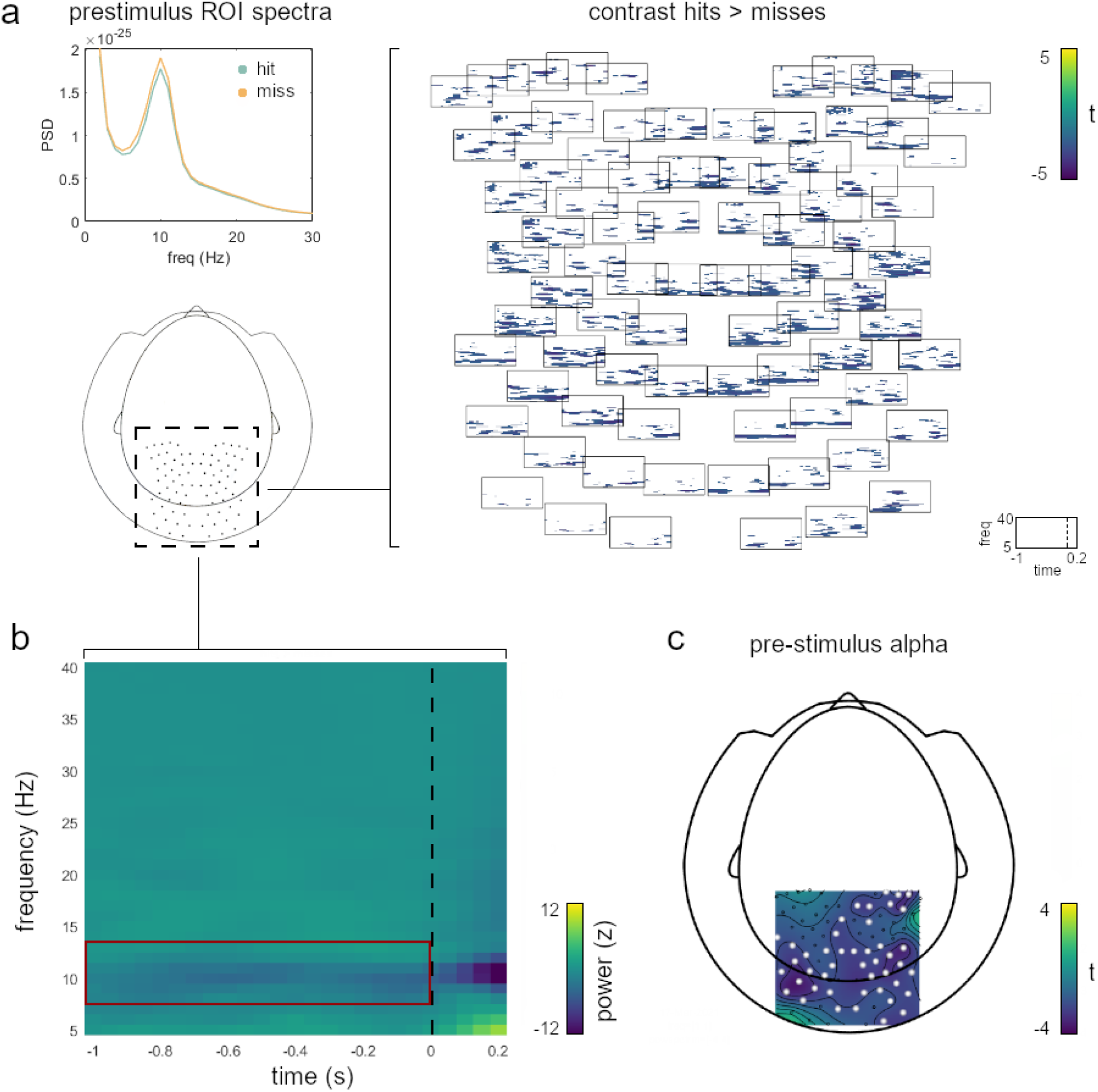
Pre- and peristimulus alpha suppression determine perceptual accuracy. **a**, Spectra for detected (hits) and undetected targets (misses) show prestimulus alpha suppression (interval [−1 0] s before target onset, top left). Right panel shows group-level significant differences (t values from cluster-corrected permutation testing) for the contrast hits > misses for single channels within the parieto-occipital ROI. Differences were computed on a frequency range of 5 - 40 Hz within a time frame of [−1 0.2] s around target onset. **b**, TFR shows power difference for the contrast hits > misses averaged across ROI sensors (same parameters as in a). Red frame indicates pre-stimulus differences in the alpha band (8 - 13 Hz). **c**, Topographic localisation of significant prestimulus alpha-band differences (as indicated in b) with significant sensors marked in white.

While these results are confirmatory rather than novel, they pose a central prerequisite for a unifying account of excitability. We suggest that respiration cyclically organises states of cortical excitability; states which in turn determine perceptual performance. Building on our findings that a) respiration cyclically modulated performance consistently across participants and b) performance was determined by phasic changes in parieto-occipital alpha power, the final questions thus concerned the overarching link between alpha power, behaviour, and respiration.

### Alpha power, behaviour, and respiration

Particularly, two key questions remained for the final analyses: First, if excitability is indeed the mediator behind respiration-locked performance changes, parieto-occipital alpha power should itself be modulated by respiration. Second, such respiration-induced changes in alpha power should contribute to the phase-dependent performance effects we saw in the PMF re-fitting, extending the global differentiation between hits and misses shown above.

As for respiration-induced changes in parieto-occipital alpha power, we replicated previous findings of phase-amplitude coupling in the alpha band ^8^. The modulation index (MI ^29^) quantifies the extent to which respiration phase modulated alpha power over parietal and occipital electrodes. Depicted in units of standard deviation of the null distribution in Fig. 4a, MI values were significant for a wide frequency range including the alpha band (tested against the 95^th^ percentile of a null distribution constructed from 200 random phase shifts, cluster-corrected for multiple comparisons at *α* = .05). This result establishes a significant respiratory modulation of the cortical parieto-occipital alpha rhythms that are known to be related to perceptual performance (cf. Fig. 3).

**Fig. 4.**
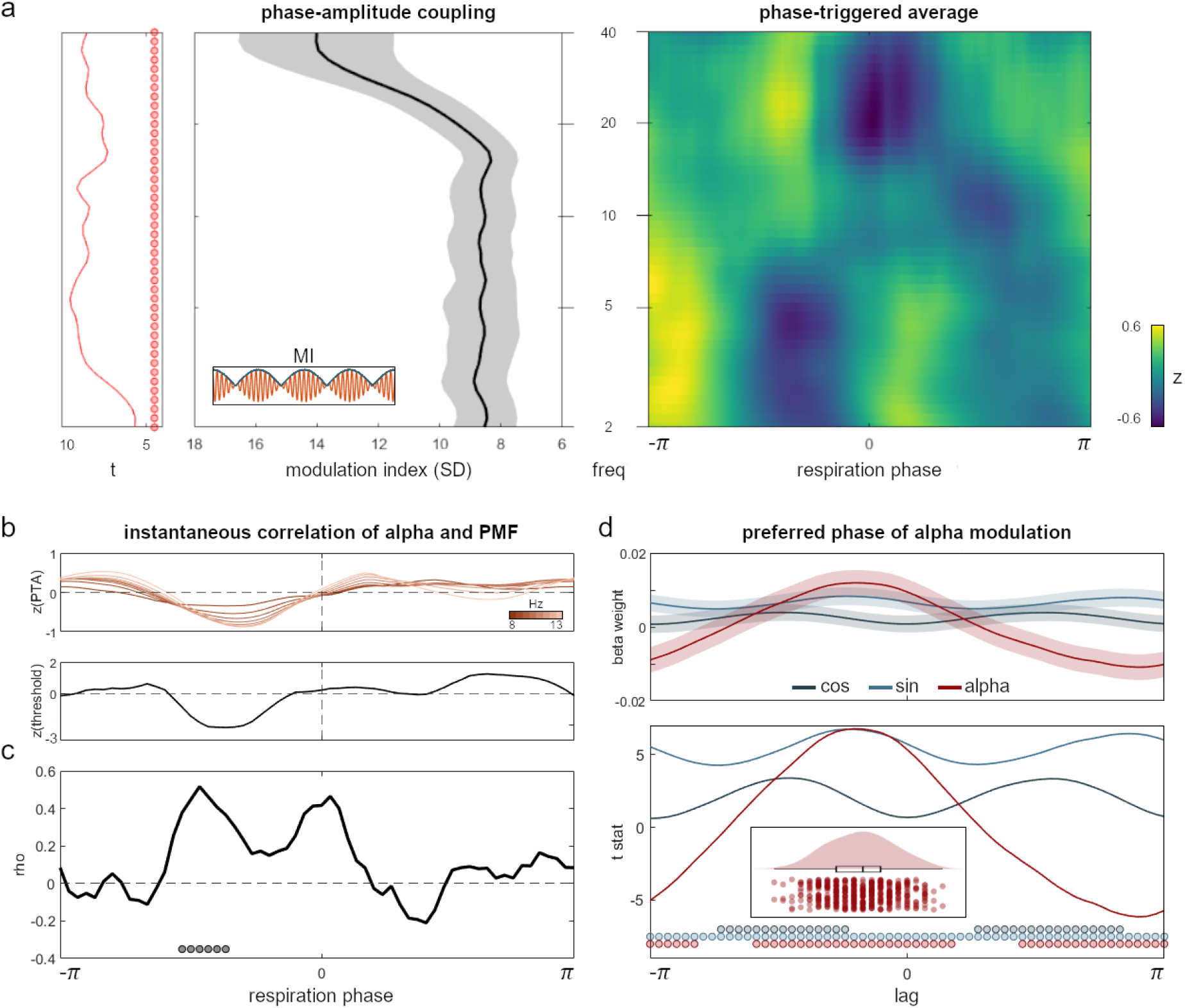
Respiration-locked alpha power modulations drive perception effects. **a**, Left panel shows modulation index as a measure of phase-amplitude coupling across frequencies and corresponding t values from cluster-permutation testing. Coupling was found significant (FDR-corrected) over the entire frequency range from 2 - 40 Hz within the parieto-occipital ROI (see Fig. 3a). Right panel shows ROI sensor-level TFR illustrating group-level average power changes over the course of one entire respiration cycle. Power was normalised within each frequency to highlight phasic changes and is shown in z values. Individual TFRs are shown in Supplementary Fig. 3. **b**, Normalised respiration-locked alpha modulation plotted against respiration-locked PMF modulation, shown here for one exemplary participant (see Supplementary Fig. 3 for individual PMF courses). c, Instantaneous group-level correlation between individual alpha and PMF threshold courses (averaged across alpha frequencies shown in b) with significant phase vectors marked by grey dots (cluster-corrected). d, LMEM beta weights (including 95% confidence intervals; top) for sine, cosine, and alpha power plus corresponding t values (bottom) shown for all phase lags between alpha and PMF courses. Significant angle vectors are marked. Individual alpha and PMF courses (as illustrated in b) were continuously shifted against each other. At each lag (ranging from −*π* to *π*), the LMEM was recomputed on the group level to obtain the preferred phase of alpha modulation effects, i.e. the lag at which alpha explained the maximum variance in PMF thresholds. Alpha effects were found to be strongest at a phase lag of − 30°, i.e. preceding PMF modulation by **π**/6. Inset shows distribution of peak lags for the alpha power coefficient from a bootstrapping approach with n = 500 iterations.

To assess the significance of alpha contributions to the interplay of respiration and performance, we set up a linear mixed effect model (LMEM) expressing the re-fitted PMF threshold values as a linear combination of sine and cosine of the respiration signal as well as parieto-occipital alpha power (averaged from 8 - 13 Hz). Significant effects were found for the sine of the respiration angle (*t*(1796) = 5.72, *p* < .001) and alpha power (*t*(1796) = 5.34, *p* < .001), providing strong support for both respiration and excitability as significant predictors of perceptual performance. Comparing the LMEM to an alternative model which only contained sine and cosine of the respiration angle confirmed the significant contribution of alpha power in explaining PMF threshold variation by means of a theoretical likelihood ratio test (*χ*^2^(1) = 28.28, *p* < .001). In other words, variation in perceptual accuracy was around 28 times more likely under the model which included parieto-occipital alpha than under one that did not.

Finally, we aimed to characterise the relationship between alpha power and PMF threshold over the respiration cycle. To this end, we computed the instantaneous (i.e. bin-wise) correlation between alpha power (averaged between 8 - 13 Hz) and PMF threshold values (exemplified in Fig. 4b). On the group level, we found the two time courses to be significantly correlated during the inspiratory phase between −90 and −60 degrees (cluster-corrected, Fig. 4c). These positive correlations corroborated our previous results showing epochs of increased performance during the inspiratory phase (Fig. 2d) as well as the perceptual benefit of suppression effects in the alpha band (Fig. 3a).

While the instantaneous correlation nicely captured interactions between alpha oscillations and PMF threshold, it was naturally computed at zero lag between the two time courses. Conceivably, however, if respiration-induced excitability changes in fact mediate performance effects, respiration-brain coupling could temporally precede respiration-induced changes in behaviour. Within the present analytical framework, this would manifest in an increased effect of alpha power at negative lags. To test this hypothesis, we combined circular shifts of the alpha power time courses with a bootstrapping approach to re-compute the LMEM described above for different lags ranging from -π to π. In short, we shifted the alpha power courses of each participant against their PMF threshold courses by a single angle bin (i.e., π/30 or 6°), re-computed the LMEM (including coefficients for sine, cosine, and alpha), and stored the beta weights and t values for each predictor. This way, we obtained a distribution of t values as a function of phase shifts which showed that the effect of alpha power peaked not at zero lag, but at a lag of −30° (Fig 4d). To test the reliability of this negative peak, we re-computed binwise LMEMs n = 500 times, each time randomly drawing 30 participants (with replacement). The bootstrapping procedure yielded a confidence interval of [−33.17 −29.25] degrees for the peak effect of alpha power, strongly suggesting that respiration-alpha coupling temporally precedes behavioural consequences.

To summarise, both respiration phase and parieto-occipital alpha power significantly explained individual variation in perceptual sensitivity (PMF threshold). While parieto-occipital alpha and PMF threshold were found to be significantly correlated during the inspiratory phase, the effect of alpha power was maximised at respiration phase lag of ~ 30°, showing that alpha modulation precedes behavioural modulation.

## Discussion

The overall picture of our findings emphatically underscores the central role of respiration in the temporal synchronisation of information sampling to phasic states of heightened neural excitability. Extending previous reports of respiration-locked performance changes, we found respiration to modulate perceptual sensitivity in a near-threshold detection task. Assessing behavioural effects over the continuous respiration signal allowed us to characterise the shape of this modulation over time, demonstrating for the first time that it appears to be more complex than a dichotomy of inspiratory vs expiratory phase. This modulatory pattern, showing the best group-level sinusoidal fit at a frequency of 1.55 times the respiration frequency, raises interesting methodological questions for future research: As the respiratory signal itself is not strictly sinusoidal, respiration-locked performance modulation is not necessarily constrained to a sinusoidal shape. Conceivably, similar to the heartbeat-evoked potential shown for the cardiac rhythm ^30^, respiratory rhythms too could trigger event-like behavioural changes that repeat with each breathing cycle. Future studies might investigate if the temporal modulation revealed here is best described by a sinusoidal function spanning the entire breathing cycle or by other models that are possibly restricted to specific time windows within a full cycle.

We then investigated parieto-occipital oscillatory power in the alpha band (8 - 13 Hz) as a proxy of neural excitability. Alpha power was significantly suppressed prior to detected vs undetected targets, highlighting the behavioural benefits of heightened prestimulus excitability. Furthermore, parieto-occipital alpha was found to be significantly modulated by respiration phase, strongly suggesting a functional coupling of respiration-locked changes in both excitability and performance. This coupling was further accentuated by a significant instantaneous correlation between alpha power and sensitivity, particularly during the inspiratory phase. Notably, the effect of alpha power on behavioural performance was strongest at a respiration phase lag of around −30°, indicating that respiration-alpha coupling temporally precedes performance changes.

### Alpha oscillations, neural excitability, and behaviour

There is overwhelming evidence for an intricate connection between alpha oscillations, excitability, and behaviour, particularly in the visual domain (see ^28^ for a recent review). In addition to alpha pacemaker cells originally shown in the animal thalamus ^31^, widespread alpha activity suggests generators to be further located in early visual ^32^ and even higher-order cortical areas ^33,34^. Excitatory input to the visual cortex is regulated by functional inhibition in a feed-forward mechanism based on alpha oscillations, effectively controlling the excitability of the neural system per se ^35,36^. The inverse relationship between alpha power and excitability is corroborated by cross-modal evidence that strong ongoing alpha oscillations entail reduced single-unit firing rates in humans ^18^ and primates ^37,38^, population-level activity such as local field potentials ^39,40^, and hemodynamic BOLD activity ^41,42^. In perceptual tasks, a transient state of lowered excitability inevitably affects behaviour, as evident from longer reaction times ^43,44^, lower confidence reports ^45,46^, and lower detection rates of near-threshold stimuli ^23,47^. Accordingly, one recent study proposed that the alpha rhythm shapes the strength of neural stimulus representations by modulating excitability ^35^. Our findings not only provide further support for the link between alpha power and excitability-related performance changes, but further suggest respiration as a key contributing factor in the ongoing search for an underlying mechanism. Such an argument for functional body-brain coupling does not contradict the proposed alpha mechanism in any way, but rather broadens the explanatory scope to unify neural and peripheral signalling.

### Excitability changes are coupled to respiration

Compared to the extensive research of alpha oscillations and their relation to neural excitability, the link between excitability and respiration is considerably less well understood. Intracranial recordings in animals have demonstrated that respiration modulates spike rates in a variety of brain regions ^14,48,49^. This characteristic was implemented in a graph model by Heck and colleagues ^50^ which provided a proof of principle that sinusoidal input and intrinsic properties of cortical networks are sufficient to potentially achieve respiration-locked modulations of high-frequency oscillations. The authors later proposed two main sources of respiration-locked neural signalling, namely the olfactory bulb (OB) and extrabulbar sources within the brainstem ^51^. As outlined in the introduction, there is broad consensus that cross-frequency coupling ^12,52^ plays a central role in translating respiratory to neural rhythms: Respiration entrains OB activity via mechanoreceptors, after which the phase of this infraslow rhythm is coupled to the amplitude of faster oscillations (see ^13,14^). As for the mechanism underlying respiration-locked excitability changes, previous work has highlighted the interplay of arterial CO_2_, tissue acidification, and adenosine. In a bidirectional pursuit of homeostasis, CO_2_ levels continuously regulate blood flow and respiration rate, which in turn determine changes in CO_2_ levels ^53^. Moreover, CO_2_ is inversely related to tissue acidity (pH), which has been linked to neural excitability ^54,55^. Lowered pH (due to increased CO_2_) causes an increase in extracellular adenosine ^56^, a neuromodulator gating synaptic transmission. Consequently, Dulla and colleagues ^17^ concluded that CO_2_-induced changes in neural excitability are caused by pH-dependent modulation of adenosine and ATP levels (also see ^57^). In our present paradigm, transient phases of heightened excitability would then be explained by decreased inhibitory influence on neural signalling within the visual cortex, leading to increased postsynaptic gain and higher detection rates. Given that the breathing act is under voluntary control, the question then becomes to what extent respiration may be actively used to synchronise information sampling with phasic states of heightened excitability.

### Towards an active account of human respiration-brain coupling

Such an active account of respiration-brain coupling is once more motivated by the animal literature. In rodents, respiration is only one of multiple rhythmic processes that constitute orofacial motor behaviours, including head and nose movements, whisking, and sniffing ^58^. Distinct respiratory nuclei in the ventral medulla have been shown to coordinate these complex motor patterns in a specific manner, in that the respiratory rhythm effectively ‘resets’ faster rhythms like whisking behaviour ^2^. According to this ‘master clock’ conceptualisation of respiration, respiratory circuits in the brain stem allow phase-locking of various sensorimotor channels through ‘snapshots’ of the orofacial environment continuously triggered by respiration onsets ^59^. In other words, multiple streams of sensory information are coordinated in a way that optimises their integration and propagation ^60^. While the concept of active sensing has thus been well established in animals ^61^, interoceptive inference in human sensation is still in its infancy. Intriguingly, theoretical modelling work ^60,62,63^ as well as empirical studies on various interoceptive signals ^64–66^ are seeking to widen our understanding of how sampling information from the external world is coordinated with bodily states. Reports on functional body-brain coupling span cyclic signals from infraslow gastric ^64^ over respiratory ^9,67^ to cardiac rhythms ^65,66^, showing that, across all time scales, sensory information is critically dependent on physiological rhythms. From a predictive processing perspective, the link between interoceptive and neural rhythms serves *predictive timing*, meaning that information sampling itself is temporally aligned with particular bodily states. Using free visual search, Galvez-Pol and colleagues ^65^ demonstrated that the timing of eye movements was closely coupled to certain phases of the cardiac cycle, with information sampling clustered during quiescent periods of the heartbeat. Functionally, the authors interpret this synchronisation to maximise the signal-to-noise balance between interoceptive and exteroceptive signals, which is precisely the mechanism we propose for the alignment with excitability states. A similar argument has prominently been made for attentional selection ^68^ which actively phase-locks neural oscillations to sensory streams in order to upregulate response gain and amplify attended stimuli ^69^. Our findings, together with mounting evidence from recent interoceptive accounts, strongly suggest that well-described accounts of hierarchically organised oscillations ^70^ are not limited to neural signals alone, but extend to even slower bodily rhythms. Recognising human respiration as active sensory selection rather than a mere bottom-up automatism offers a promising framework to explain how respiration-locked changes in neural signalling benefit action, perception, and cognition.

## Acknowledgements

We would like to thank Karin Wilken, Ute Trompeter, and Hildegard Deitermann for their invaluable assistance during data collection. This work was supported by the Interdisciplinary Center for Clinical Research (IZKF) of the medical faculty of Münster (Gro3/001/19). NAB (BU2400/9-1) and JG (GR 2024/5-1) were further supported by the DFG.

## Methods

### Participants

Thirty volunteers (16 female, age 25.1 ± 2.7 years [mean ± SD]) participated in the study. All participants had (corrected-to) normal vision, denied having any respiratory or neurological disease, and gave written informed consent prior to all experimental procedures. The study was approved by the local ethics committee of the University of Muenster and complied with the Declaration of Helsinki.

### Procedure

Participants were seated upright in a magnetically shielded room while we simultaneously recorded respiration and MEG data. MEG data were acquired using a 275 channel whole-head system (OMEGA 275, VSM Medtech Ltd., Vancouver, Canada) and continuously recorded at a sampling frequency of 600 Hz. To minimise head movement, participants’ heads were stabilised with cotton pads inside the MEG helmet. Data were acquired across six runs with intermediate self-paced breaks. The length of each run was dependent on individual response speed (452 ± 28 s; M ± SD). Participants were to breathe automatically through their nose while respiration was recorded as thoracic circumference by means of a respiration belt transducer (BIOPAC Systems, Inc., Goleta, United States) placed around their chest. To characterise individual rates of natural breathing, we computed time-frequency representations of single-run respiration time courses for each participant. To this end, the continuous respiration signal was segmented into 20 s segments with 50% overlap and subsequently detrended. Power spectra (range 0 to 2 Hz in .025 Hz increments, single Hanning taper, zero-padded to 40s segments) were computed in Fieldtrip ^71^ and subsequently averaged across runs within each participant. On the group level, the dominant breathing frequency was found to be at 0.26 ± 0.08 Hz (M ± SD), corresponding to an average of 15.8 breaths per minute.

### Task

Participants performed a spatial detection task in which they were to fixate on a cross (0.4 degrees in diameter) in the centre of the screen presented against a black background. Following this fixation period (jittered between 1200 and 3500 ms), a small Gabor patch (0.3 degrees in diameter) was presented for 50 ms in a marked circular area (3.5 degrees in diameter) 10 degrees to the left or the right side of the fixation cross. In addition to left or right side targets, one third of all trials were catch trials where no target was presented at all. After a delay of 500 ms, a question mark in the centre of the screen prompted participants to give their response: Using a 3-button response box in their right hand, participants reported whether they saw a target on the left (index finger) or the right side (middle finger) or no target at all (thumb). Once the report was registered, a new trial started (again with a fixation period). Participants were instructed to keep their eyes on the centre cross at all times and encouraged to report left or right targets even when they were not entirely certain.

For each trial, target contrast was adapted by a QUEST staircase ^72^ aimed at individual hit rates of about 60 %. Each of the six experimental runs contained a total of 720 trials (240 left target, right target, and catch trials, respectively).

### Behavioural analyses

Behavioural data were preprocessed and analysed using R (R Core Team, 2014) and Matlab (The Mathworks, Inc., Natick, United States). To account for the initiation of the QUEST procedure, the first 20 trials of each run were discarded. All behavioural analyses were focussed on trials with an actual target present (n = 480) which were subsequently classified into hits (i.e., detected targets) and misses (undetected targets). Individual hit rates (HR) were computed as n_hits_ / (n_hits_ + n_misses_).

To obtain the respiration phase angles corresponding to single trials, we used Matlab’s *findpeaks* function to identify time points of peak inspiration (peaks) and expiration (troughs) in the normalised respiration time course. Phase angles were linearly interpolated from trough to peak (-π to 0) and peak to trough (0 to π) in order to yield respiration cycles centred around mean inspiration. For each trial, we thus obtained the respiration angle at which the target had been presented.

Using the *gcmi* toolbox for Matlab ^73^, we computed single-trial mutual information on the participant level for two sets of factors: First, we considered a discrete *detection* factor (target detected/undetected) and a continuous *contrast* factor of Copula-normalised target contrast (as determined by the QUEST procedure). The Copula transform is based on rank ordering the individual target contrast vectors (see ^73^ for details). In a second step, we added two further continuous *sine* and *cosine* factors of respiration phase. These computations yielded two mutual information values for each participant, with and without respiration phase, respectively. To determine whether adding respiration phase significantly enhanced mutual information on the group level, we tested individual differences of the two mutual information values against zero with a Wilcoxon signed rank test.

### Respiration phase-dependent PMF fitting

For a more sensitive measure of respiration-locked changes in perceptual accuracy, we combined the moving window approach outlined above with a robust Bayesian inference analysis for the psychometric function. For each participant, we first fitted the psychometric function (with a cumulative Gaussian distribution as the sigmoid) to all target trials using a three-alternative forced choice model (guessing rate fixed at .33) from the *Psignifit* toolbox for Matlab ^74^. We then used the identified threshold and width (i.e. slope) from the overall fit as priors for an iterative refitting of the PMF on subsets of trials obtained from the moving window: Again moving along the respiration cycle in increments of Δω = π/30, we fitted the PMF to the trials presented at a respiration angle of ω ± π/10. This way, we extracted respiration phase-dependent estimates of PMF threshold for each participant (see Fig. 2) which were subsequently z-scored. Finally, the normalised threshold variations were averaged across subjects to obtain the grand average phase-dependent threshold modulation at the population level. Next, we fitted a sinusoid function in the following form:

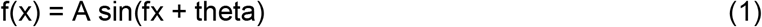

using the *Curve Fitting* toolbox for Matlab. The procedure yielded the adjusted R^2^ as a criterion to correct the goodness of fit for the number of parameters used in our model (three, in our case: amplitude, frequency, and phase of the sinusoidal model). For significance testing at the population level, we performed a nonparametric permutation test ^75,76^ by randomising the phase angles at the single trial level separately for each participant. In repeating the previously described analysis 1000 times on these randomised data, we constructed a null distribution of adjusted R^2^ values. This approach allows to reject the null hypothesis (in our case, no sinusoidal modulation of threshold as a function of phase angle) when the adjusted R^2^ obtained on the model fitting exceeds the 95^th^ percentile of the adjusted R^2^ null distribution. In order to characterise phase consistency between subjects at the best fitted frequency, we first computed the fast Fourier transform (FFT) of individual threshold variation series, after zero padding in order to increase our frequency resolution. Then we selected for each participant the phase angle of the vector in the complex plane closest to the fitted frequency and tested for angle uniformity across subjects with the Rayleigh test using the *CircStat* toolbox for Matlab ^77^.

### MEG preprocessing and segmentation

All MEG and respiratory data preprocessing was done in Fieldtrip for Matlab. Prior to statistical comparisons, we adapted the synthetic gradiometer order to the third order for better MEG noise balancing (*ft_denoise_synthetic*). Power line artefacts were removed using a discrete Fourier transform (DFT) filter on the line frequency of 50 Hz and all its harmonics (including spectrum interpolation; *dftfilter* argument in *ft_preprocessing*). Next, we applied independent component analysis (ICA) on the filtered data to capture eye blinks and cardiac artefacts (*ft_componentanalysis* with 32 extracted components). On average, artefacts were identified in 2.53 ± 0.73 components (M ± SD) per participant and removed from the data. Finally, continuous MEG data were segmented to a [−1.25 0.75] s interval around target onset and resampled to 300 Hz.

Likewise, respiration time courses were resampled to 300 Hz and z-scored to yield normalised respiration traces for each participant and each experimental run.

### Head movement correction

In order to rule out head movement as a potential confound in our analyses, we used a correction method established by Stolk and colleagues ^78^. This method uses the accurate online head movement tracking that is performed by our acquisition system during MEG recordings. This leads to six continuous signals (temporally aligned with the MEG signal) that represent the x, y, and z coordinates of the head centre (H_x_, H_y_, H_z_) and the three rotation angles (H_ψ_, H_*υ*_, H_φ_) that together fully describe the head movement. We constructed a regression model comprising these six ‘raw’ signals as well as their derivatives and, from these 12 signals, the first, second, and third-order non-linear regressors to compute a total of 36 head movement-related regression weights (using a third-order polynomial fit to remove slow drifts). This regression analysis was performed on the power spectra of single-sensor time courses, removing signal components that can be explained by translation or rotation of the head with respect to the MEG sensors.

### MEG time-frequency analyses

Parieto-occipital time-frequency representations (TFRs) of hit and miss trials were computed using a discrete prolate spheroidal sequences (DPSS) multitaper on a frequency range from 5 to 40 Hz (*mtmconvol* argument for *ft_freqanalysis*). We applied a 2 Hz smoothing kernel to frequencies below 30 Hz and a 5 Hz smoothing kernel to frequencies above 30 Hz. As hits were slightly more frequent than misses (recall that the QUEST staircase aimed to fix performance at a hit rate of .60), we randomly sampled hit trials to match the number of misses, thus avoiding sampling bias. Time-frequency representations were computed for a moving window of 500 ms (50 ms increments) over the entire time frame of [−1.25 0.75] s around target onset. TFRs of hits and misses were separately averaged across participants and collapsed over the time dimension (range [−1 0] s) to obtain pre-stimulus power spectra (see Fig. 4a).

Again, we used non-parametric permutation testing to determine statistical significance of group-level differences between TFRs of hits and misses. Individual TFRs were represented in matrices of 82 ROI channels x 36 frequencies x 25 time points. Differences across participants were assessed by means of dependent-samples t-tests while correcting for multiple comparisons (across channels, frequencies, and samples) with a Monte Carlo-based cluster permutation approach (5000 randomisation iterations, α = .05).

### Computation of global field power

For the computation of global field power, the time courses of 80 MEG channels within our parieto-occipital ROI were individually subjected to a continuous wavelet transform using a Morlet wavelet with a Fourier-based algorithm for 64 frequencies (ranging from 0.5 to 40 Hz). The Fourier transform of our wavelet is defined as

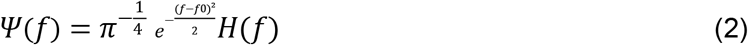

where H(f) is the Heaviside function and f0 is the centre frequency in radians/sample. Next, we computed the absolute amplitude envelope of this complex-valued data, smoothed it with a 300ms moving average, and averaged these amplitude values across channels and experimental runs.

### Computation of modulation index and phase-triggered average

The modulation index (MI) quantifies cross-frequency coupling and specifically phase-amplitude coupling. Here, it was used to study to what extent the amplitude of brain oscillations at different frequencies were modulated by respiration phase. To this end, the instantaneous phase of the respiration time course was computed with the Hilbert transform. MI computation was then based on the average oscillatory amplitude (ranging from 2 - 40 Hz) at 20 different phases of the respiratory cycle (centred around peak inspiration). Any significant modulation (i.e. deviation from a uniform distribution) is quantified by the entropy of this distribution. To account for frequency-dependant biases, we followed previously validated approaches ^7^ and computed 200 surrogate MIs using random shifts of respiratory phase time series with concatenation across the edges. The normalised MI was computed by subtracting, for each frequency, the mean of all surrogate MIs and dividing by their standard deviation leading to MI values in units of standard deviation of the surrogate distribution. The computation resulted in normalised MI values for each channel, frequency, and participant. Group-level significance of these normalised MI values was determined by means of cluster-based permutation testing using *ft_freqstatistics* in Fieldtrip. Specifically, we conducted a series of one-tailed t-tests of individual MI values at each frequency against the 95th percentile of the null distribution from the 200 surrogate MI values. T values were then thresholded at p = .05 and adjacent significant data points were defined as clusters. Using 5000 randomisation iterations and the cluster sum criterion, the original cluster statistics were compared to the histogram of the randomised null statistics. Clusters in the original data were deemed significant when they yielded a larger test statistic than 95% of the randomised null data.

To assess oscillatory modulation over time, the phase-triggered average (PTA) was computed from the smoothed, band-specific amplitude envelopes averaged across the 80 ROI sensors. Time points of peak inhalation were detected from the respiration phase angle time series using Matlab’s *findpeaks* function. For each peak in the respiration signal, global field power across all 64 frequencies was averaged within a time window of ± 1000 samples centred around peak inhalation. The resulting 64 frequencies x 2000 samples matrix was finally z-scored across the time dimension, leading to normalised oscillatory power within the ROI phase-locked to the respiration signal. This analysis is equivalent to a wavelet-based time-frequency analysis. Computations were done separately for the 6 experimental runs and subsequently averaged across runs and participants.

### Instantaneous correlation

We investigated the presence of an instantaneous, positive group-level correlation between variations in prestimulus alpha power and variations in the PMF threshold, both as a function of the respiration phase angle. First, we averaged the alpha power over the range between 8-13 Hz. Then, for each phase angle bin, we correlated the alpha power with the corresponding threshold values. This yielded a course of Pearson *r* and corresponding p values over the respiration cycle. In order to correct for multiple comparisons, we applied a cluster permutation approach as follows: We first detected the significant clusters in our data and obtained the cluster masses by summing together the *r* values of the contiguous significant points (p < .05). Then we reiteratively shuffled (N=10000) the participant order in both the matrices of alpha power and thresholds. For each iteration, we stored the highest cluster mass value, yielding a surrogate distribution of cluster masses. We rejected H0 for each cluster mass in the real data that exceeded the 95^th^ percentile (α = .05, one-tailed) of the surrogate distribution of cluster masses.

### Linear mixed effect modelling

We employed linear mixed effect modelling (LMEM) to investigate the relationship between respiration phase, parieto-occipital alpha power, and perceptual accuracy. LMEM models a response variable (in our case, the PMF threshold value) as a linear combination of fixed effects shared across the population (phase angle, alpha power) and participant-specific random effects (i.e modulatory variation between participants). We specified two LMEMs: The first model was set up to predict the PMF threshold as a function of respiration angle (with separate sine and cosine contributions):

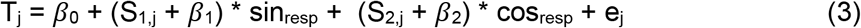

For participant j, the PMF threshold value is expressed as a combination of the intercept (*β*_0_), the fixed effects of sine and cosine of the respiration angle (*β*_1_, *β*_2_), and an error term (e_j_ ~ N(0,σ^2^)). Since we had observed considerable between-participant variation of respiration-locked PMF changes during the previous analysis (see above), we specified the current LMEM to include random slopes (S_1,j_, S_2,j_). This accounts for the fact that a potential fixed effect of respiration phase does not necessarily have to modulate PMF threshold identically across the entire cohort. An analogous approach was used in a second model that further included the fixed effect of alpha power:

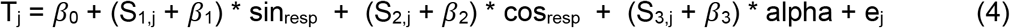

We used Matlab’s *compare* function to test whether including alpha power in the model would significantly increase the LMEM fit (while simultaneously accounting for the addition of a third fixed effect) by means of a theoretical likelihood ratio test.

Next, we attempted to characterise potential phase shifts between the modulatory effects of respiration on PMF thresholds and alpha power. To this end, for lags ranging from -π to π, we shifted the alpha power course of each participant against their PMF threshold course by a single angle bin (i.e., π/30 or 6°), re-computed the group-level LMEM (including coefficients for sine, cosine, and alpha), and stored the beta weights and t values for each predictor. This way, we obtained a distribution of t values as a function of phase shifts between alpha and PMF time courses.

To quantify the reliability of the ideal phase lag we obtained at around −30° (Fig. 4d), we combined the previous permutation procedure with a bootstrapping approach: For n = 500 iterations, we randomly selected 30 participants (with replacement) and re-computed binwise LMEMs as described above.

### Data availability

The anonymised data supporting the findings of this study are openly available from on the Open Science Framework (https://osf.io/ajuzh/).

