## Supplementary Figures for "Respiration aligns perception with neural excitability"

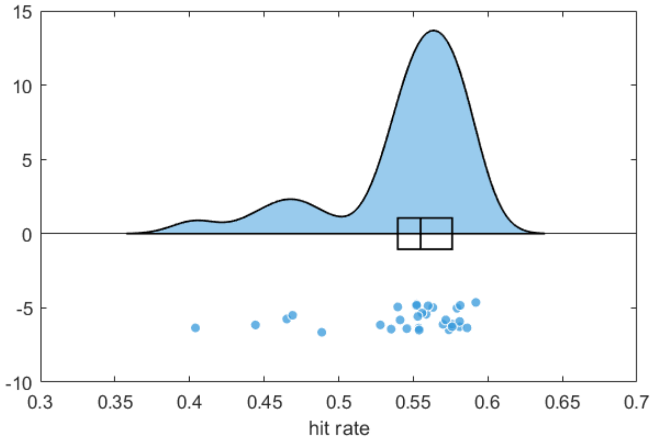


**Supplementary Fig. 1.** Group-level distribution of individual hit rates across all trials.

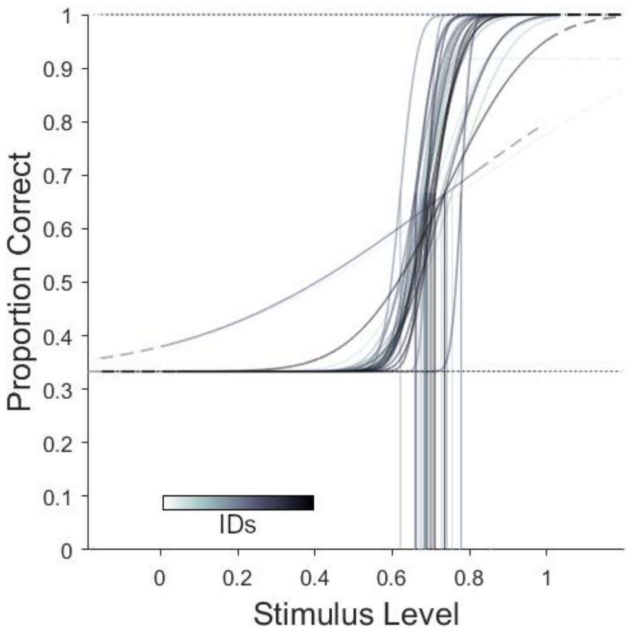


**Supplementary Fig. 2.** Individual psychometric functions (PMF), fitted on the full data set. For each participant, PMF changes across the respiration cycle were assessed by subsequently fixing all parameters except the PMF threshold (vertical lines) and re-fitting the PMF on subsets of trials falling into one of n = 60 respiration phase bins (exemplified in Fig. 2b of the main manuscript).

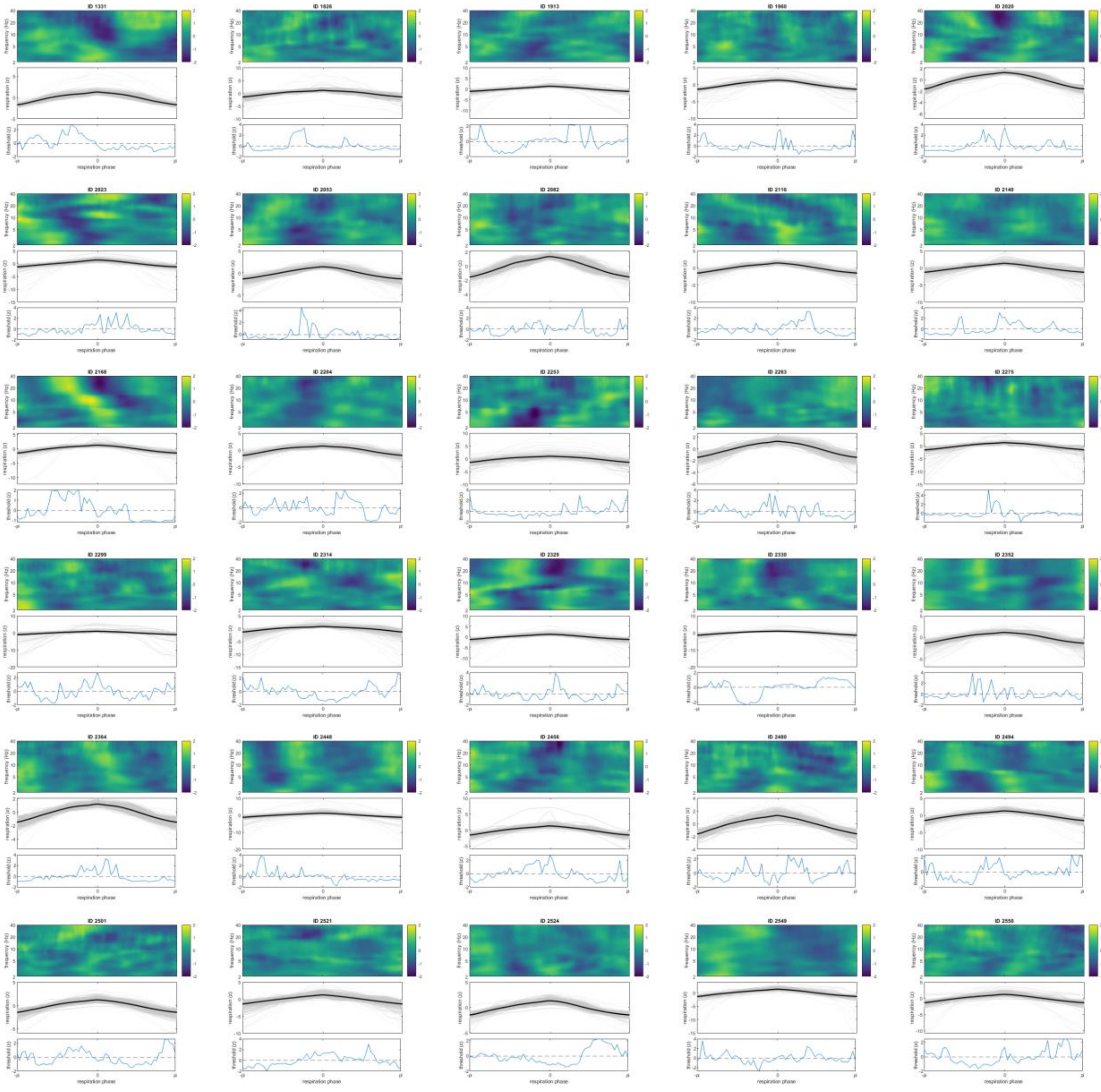

**Supplementary Fig. 3.** Individual phase-triggered averages (top), normalised respiration time courses (plus mean respiration time course in bold; middle), and normalised PMF threshold over the respiration cycle (bottom).
